# Addressing complex autofluorescence signatures in solid tissue samples to enhance full spectrum flow cytometry of non-immune cells

**DOI:** 10.64898/2025.12.19.695385

**Authors:** Christina Gkantsinikoudi, Manuela Terranova-Barberio, Neil P Dufton

## Abstract

FSFC is an emerging technology that can greatly enhance our understanding of the single-cell proteomic landscape. However, its application to cells derived from solid tissues has been hampered by their complex autofluorescence signatures and lack of optimized tools for non-immune cells. Here, we present a protocol and discuss key controls that minimize the impact of unmixing errors enabling us to resolve multiple EC subpopulations isolated from different tissues in models of chronic tissue injury.

**Research Topic(s):** Vascular biology, cell heterogeneity, full spectrum flow cytometry

**Graphical abstract:** 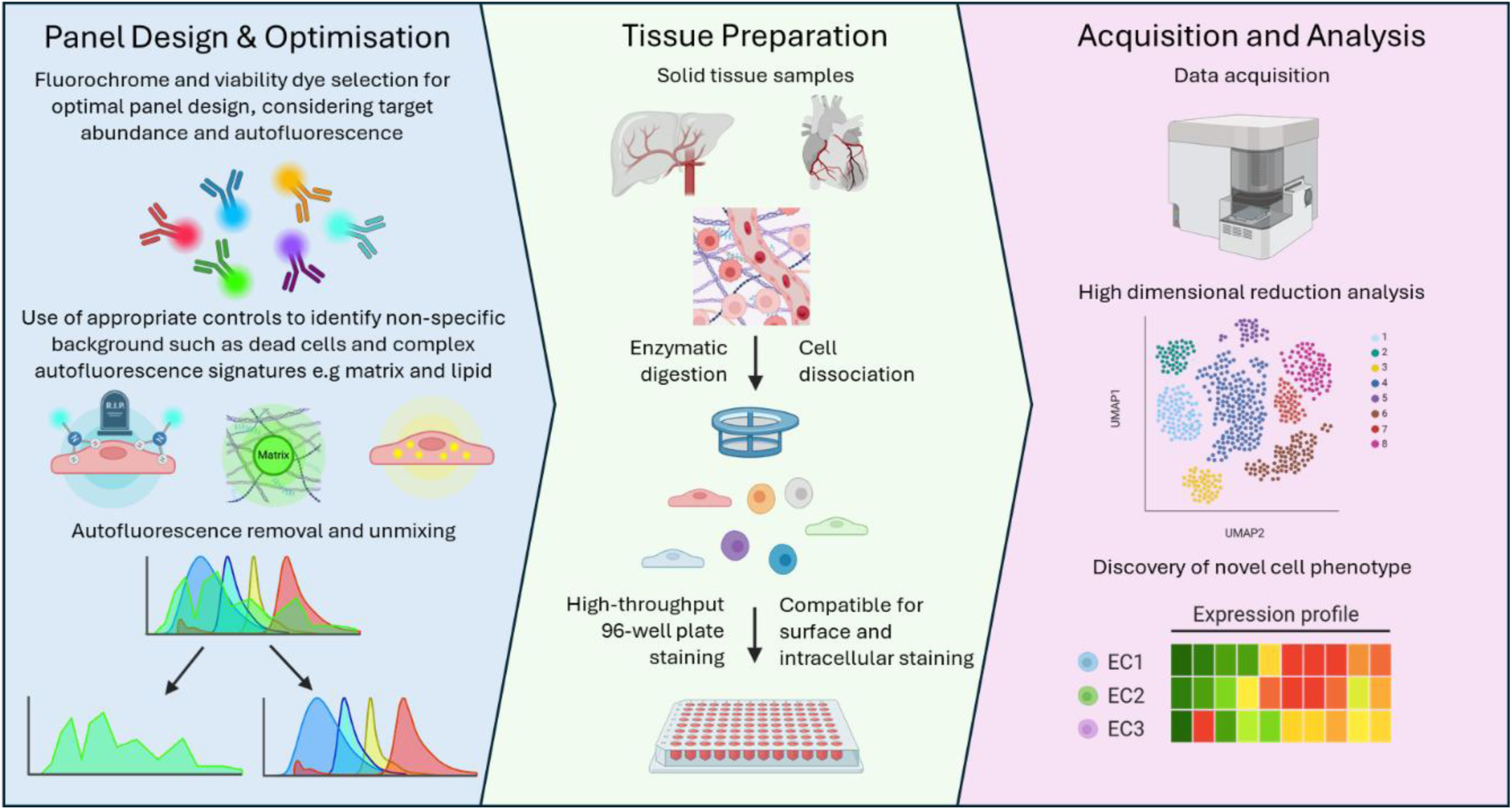

**Highlights:** Optimisation of a FSFC panel to enable in-depth phenotyping of tissue- and model-specific endothelial subpopulations from solid tissues.

Discussion of appropriate controls to minimize the impact of tissue autofluorescence and enhance the signal-to-noise ratio for cell phenotyping in complex models of inflammation and fibrosis.

Trajectory analysis to track cellular plasticity over time.

Application of full spectrum cell sorting to isolate rare endothelial subpopulations with complex phenotypes.

## Introduction

Phenotyping of single cells has provided critical insights into cellular heterogeneity and function with major implications for developing novel targeted therapies. Single-cell RNA sequencing (scRNAseq) has become the gold standard approach for characterising tissue- and disease-specific cellular heterogeneity. However, reliance on transcriptomic data has key limitations: RNA abundance does not always parallel protein expression, and information regarding post-transcriptional or post-translational modifications, as well as cell-surface protein-mediated interactions, are not captured ^1^.

Full spectrum flow cytometry (FSFC) is an emerging technology that addresses some of these shortcomings by enabling high-dimensional, proteomic, single-cell phenotyping. By measuring protein expression directly, FSFC complements scRNAseq, offering a more comprehensive view of cellular phenotypes. Currently, FSFC is predominantly used for immuno-oncology ^2^. Its application to non-immune cell populations and heterogenous solid tissue samples presents additional challenges, as intricate autofluorescence (AF) signatures complicate panel design and spectral unmixing ^3^.

Endothelial cells (ECs) exemplify a challenging cell population for FSFC. Most studies of EC heterogeneity in health and disease have relied on scRNAseq-based phenotypic assessment ^4, 5, 6^, while single-cell proteomic analysis remains limited, in part due to their challenging AF signatures ^7^. This issue is amplified in steatotic or fibrotic tissues, where excessive lipids and extracellular matrix increase AF ^8, 9^.

To address these challenges, we developed and optimized a FSFC protocol for phenotyping ECs from murine liver and heart samples. Here, we detail critical considerations, including the selection of an appropriate viability dye, fluorochrome optimisation and strategies for efficient AF extraction, emphasising the importance of tissue-, sex- and model-specific controls to account for different sources of AF. This protocol also describes a downstream analytical pipeline for dimensionality reduction, cluster visualisation and trajectory analysis of diverse endothelial subpopulations. Furthermore, we demonstrate the successful isolation of rare EC subpopulations through full spectrum sorting.

This study provides a versatile framework for applying FSFC to diverse solid tissues and non-immune cell types. Furthermore, it highlights the ability of FSFC to aid the discovery of rare cell populations, describe phenotypic changes and cell plasticity^10^, and has the potential to identify new therapeutic targets.

## Results

### Panel design

When determining the size and spectral range of a panel, it is important to consider the capabilities of the equipment, such as the number of lasers and detectors available, which will dictate the range of fluorochromes that are compatible. Availability of more detectors can lead to lower panel complexity scores even for the exact same fluorochromes, thereby reducing unmixing errors. The panel we designed was for the 4 laser Cytek® Aurora (UV/V/B/R, 54 + 3 channel) platform. To calculate the fluorochrome similarity and the complexity index of the panel, the Spectrum Viewer tool (available through the Cytek® Cloud) can be used. We show the similarity score for the 14 fluorochromes in our panel with similarity ranging from 0-0.94 (Fig. 1A); Cytek® recommend that similarity can be tolerated up to 0.98 within a panel. The complexity of the panel is a measure of the spectral uniqueness of its fluorochromes, with lower values predictive of a simpler unmixing process. The stain reduction index matrix (Fig. 1B) is used to quantify how the stain index of each fluorochrome is reduced due to the presence of the other fluorochromes within the panel. Finally, the spillover matrix (Fig. 1C) identifies potential spillover of the emission wavelengths of each fluorochrome into other channels. Together these tools act as a guide for the selection of fluorochromes with unique signatures as well as highlight potential incompatible fluorochrome combinations. However, all combinations require experimental optimisation including antibody titrations and assessment of the impact of complex autofluorescence signatures to minimize unmixing errors which we will address in more detail.

**Figure 1:**
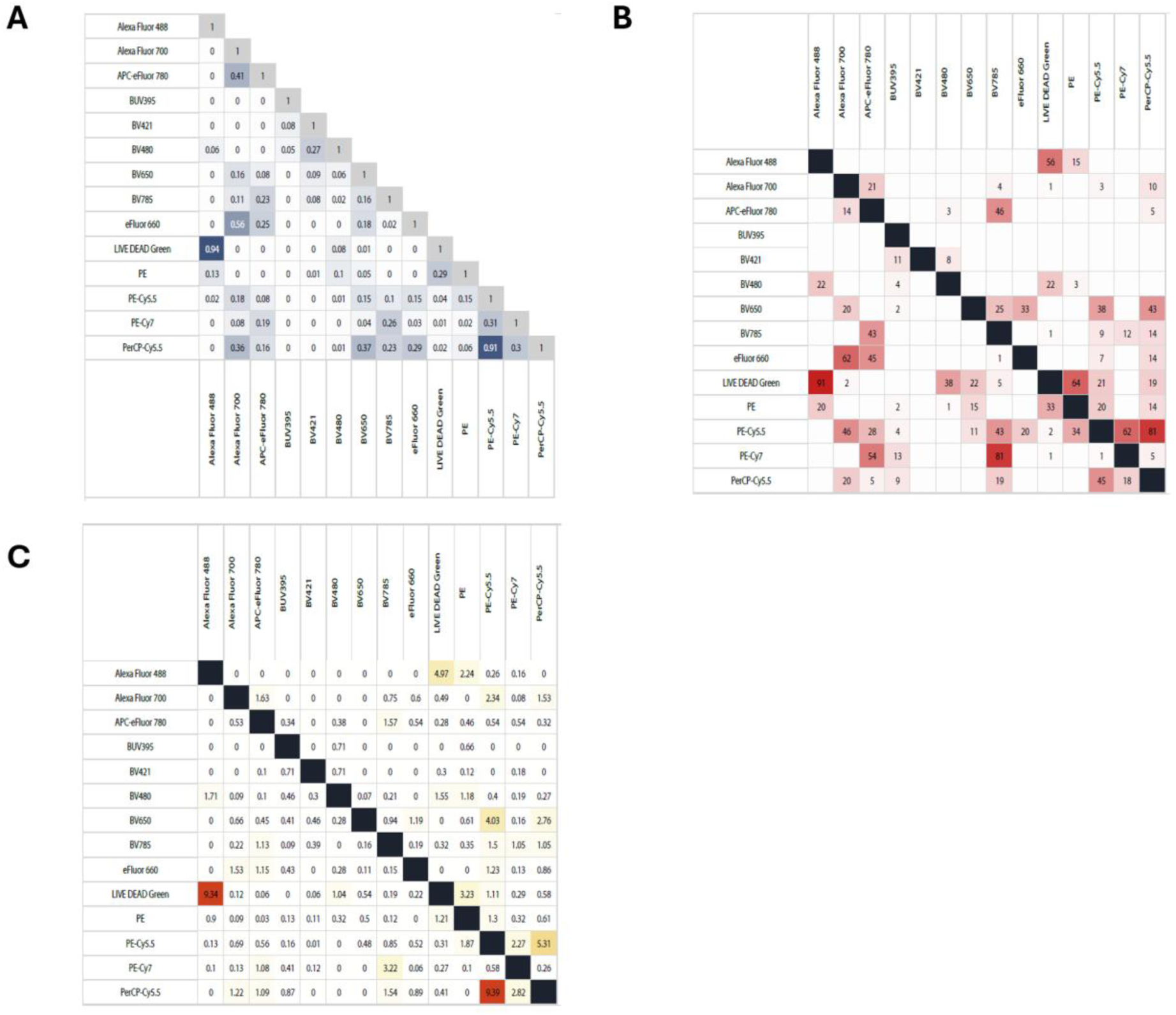
Properties of the 14-colour endothelial cell panel. The 14 fluorochromes used in the endothelial cell antibody panel were assessed using the Cytek® Spectral Viewer tool to generate a (A) complexity index, (B) stain reduction index and (C) spillover spread matrix for the panel.

To identify specific cell types within whole tissue samples it is useful to define inclusion and exclusion markers to enhance the analysis of a chosen population. To build a comprehensive EC panel, we consulted reports that used several different experimental approaches to identify and study different EC subpopulations. These approaches included lineage tracing studies ^11, 12^, transcriptomic studies ^13, 14^ and EC assessment by immunohistochemistry (IHC) ^15, 16^. Our own assessment using IHC allowed us to visualize the selectivity of different markers for specific EC subpopulations, exemplified in Figure 2 by the expression of thrombomodulin (TM) and endomucin (EMCN). Both markers are EC-specific and highly zonated. Within the hepatic vasculature (Fig. 2A), low expression can be observed in peri-portal (PP) regions, while high expression is noted around the central veins (CV) (Fig. 2B; explored in more detail in Gkantsinikoudi *et al.,* 2026*). In the heart, the same markers demonstrate ventricular zonation (Fig. 2C), displaying higher expression in endocardial EC and epicardial regions with lower expression in microvascular EC of the mid-ventricular regions (Fig. 2D). Based on this approach, 6 EC identity markers were chosen for this FSFC panel due to their EC-restricted expression profiles and their ability to identify different subpopulations within both the liver and heart tissue. These EC markers were combined with 3 markers of EC activation (ICAM1, LY6A and CD34) and 3 markers of mesenchymal transition (transgelin (TAGLN), THY1.2, and platelet-derived growth factor receptor α (PDGFRα)) allowing us to study the dynamic changes in EC phenotypes and induction of endothelial-to-mesenchymal transition (EndMT) in models of fibrosis^10^. The pan-leukocyte marker CD45 was also included in our panel to exclude CD31^+^ immune cells, while a viability dye was also used to exclude dying or dead cells from the analysis. The final gating strategy used to identify the total population of endothelial cells is presented in Fig. 3A-D.

**Figure 2:**
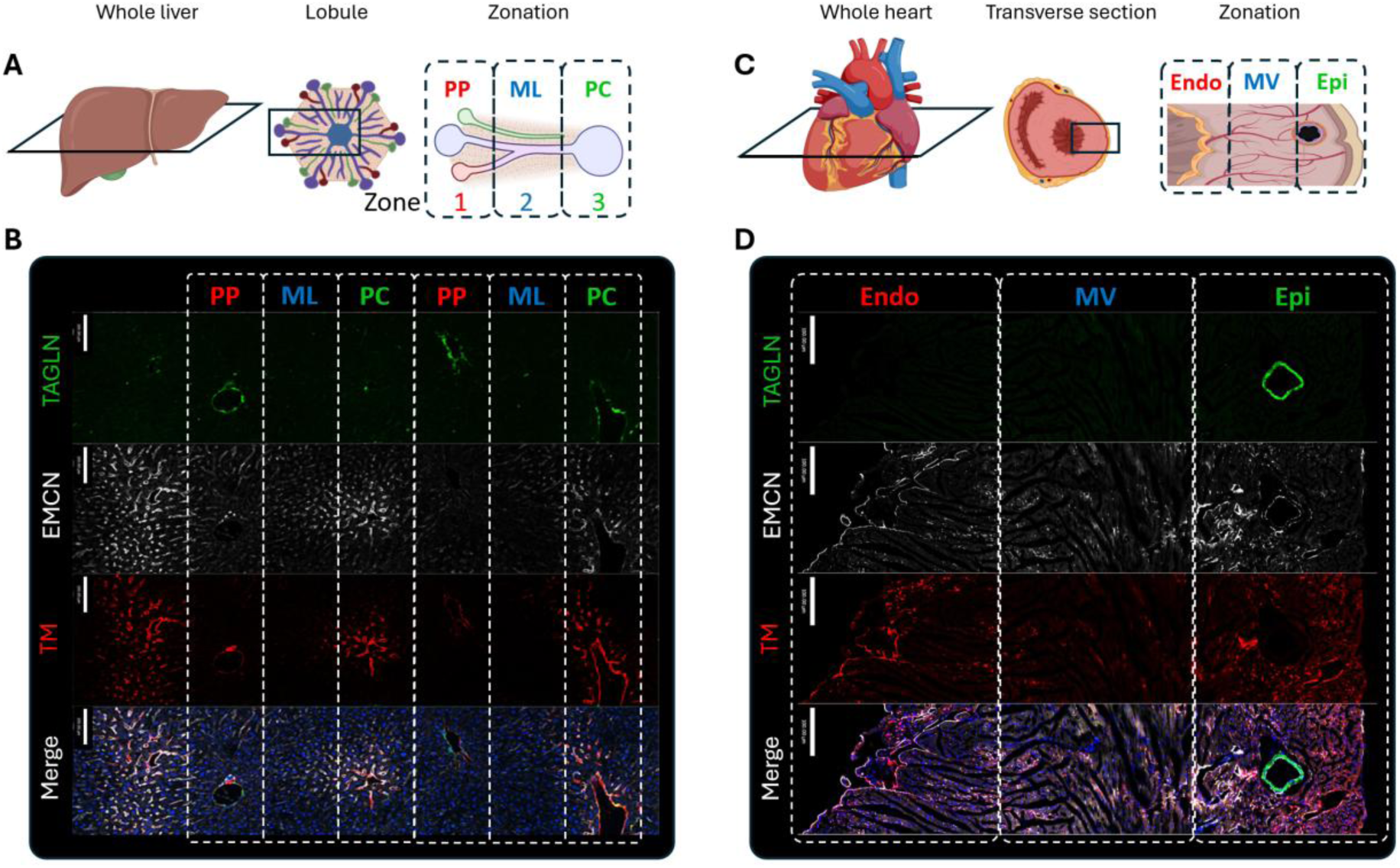
Zonated expression of endothelial markers in the liver and heart. (A) Schematic of a liver cross section. The liver is organized into hexagonal lobules that consist of three distinct zones, termed peri-portal (PP; Zone 1), mid-lobular (ML; Zone 2) and peri-central (PC; Zone 3) zones. (B) Representative confocal image of a liver section stained for transgelin (TAGLN, green; smooth muscle cells/mesenchymal cells), endomucin (EMCN, white; endothelial cells), thrombomodulin (TM, red; endothelial cells) and Hoechst (blue; nuclei). Scale bars= 100μm. (C) Schematic of a transverse cross section of the heart. The left ventricle is presented from the endocardium (Endo; left) through the mid-ventricle (MV) to the epicardium (Epi; right). (D) Representative confocal image of a heart section stained for transgelin (TAGLN, green; smooth muscle cells surrounding the coronary artery), endomucin (EMCN, white; endothelial cells), thrombomodulin (TM, red; endothelial cells) and Hoechst (blue; nuclei). Scale bars= 100μm.

**Figure 3:**
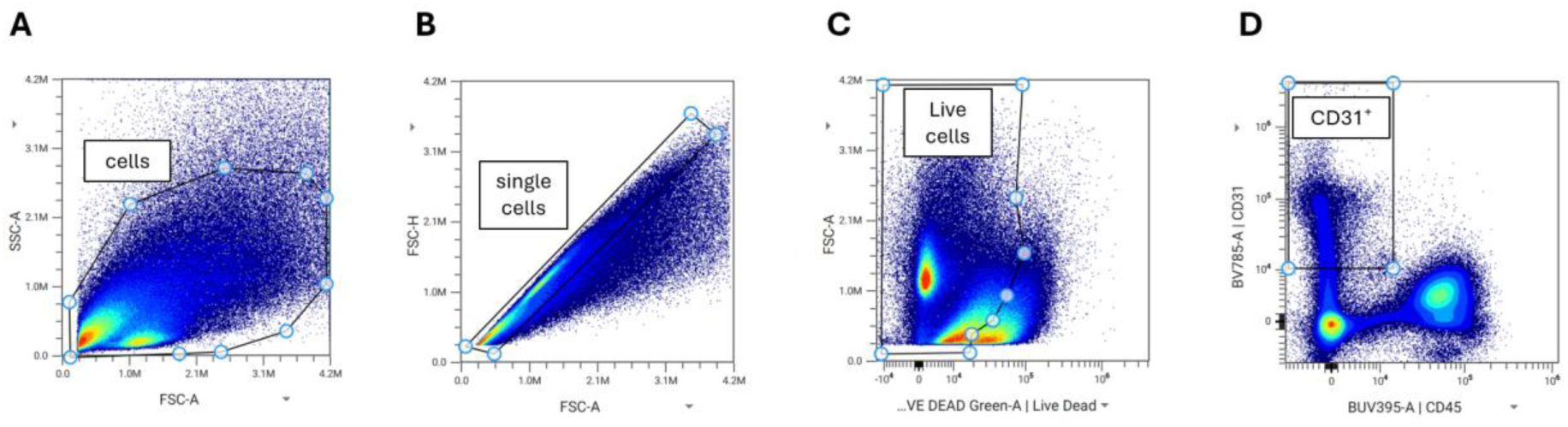
Gating strategy for the identification of endothelial cells. (A) The total cell population was identified using the FSC-A vs SSC-A plot. (B) Doublets were excluded based on the FSC-A vs FSC-H plot. (C) Live cells were selected using the LIVE/DEAD Green Fixable viability dye. (D) The total population of endothelial cells was selected based on positive signal for CD31 and negative signal for CD45 (CD31^+^CD45^-^ cells). All EC subpopulations were identified in CD31^+^CD45^-^ gates.

### Optimisation of the viability dye and antibody dilutions

#### Viability dye selection

The selection of a suitable viability dye is the crucial first step in optimizing a successful panel. Two of the most common and spectrally diverse types of viability dyes are DNA-binding (e.g. DAPI or Hoechst) and amine-reactive dyes (e.g. LIVE/DEAD Fixable Viability Dyes (Invitrogen), Zombie Fixable Viability Dyes (Biolegend)). We found that using a DNA-binding dye was not the most suitable approach for our panel, as due to the fixation and permeabilisation step of our protocol, DNA staining of live cells was evident, even after optimisation efforts. Therefore, we assessed the staining profiles of three LIVE/DEAD Fixable Dye reagents to identify a potentially compatible fluorochrome to include in the panel. There are striking differences in the cellular distribution profiles between the three fluorochromes tested (Fig. 4A-C) with Live/Dead Green chosen for the clearest segregation of viable cells. This was contrary to our initial assessment that NIR would be the most suitable dye due its emission wavelength at a low-AF zone of the spectrum. However, one of the chemical compounds used to generate the liver injury mouse model resulted in high AF in that region. This highlights the importance to consider model-specific sources of AF when designing a FSFC panel.

**Figure 4:**
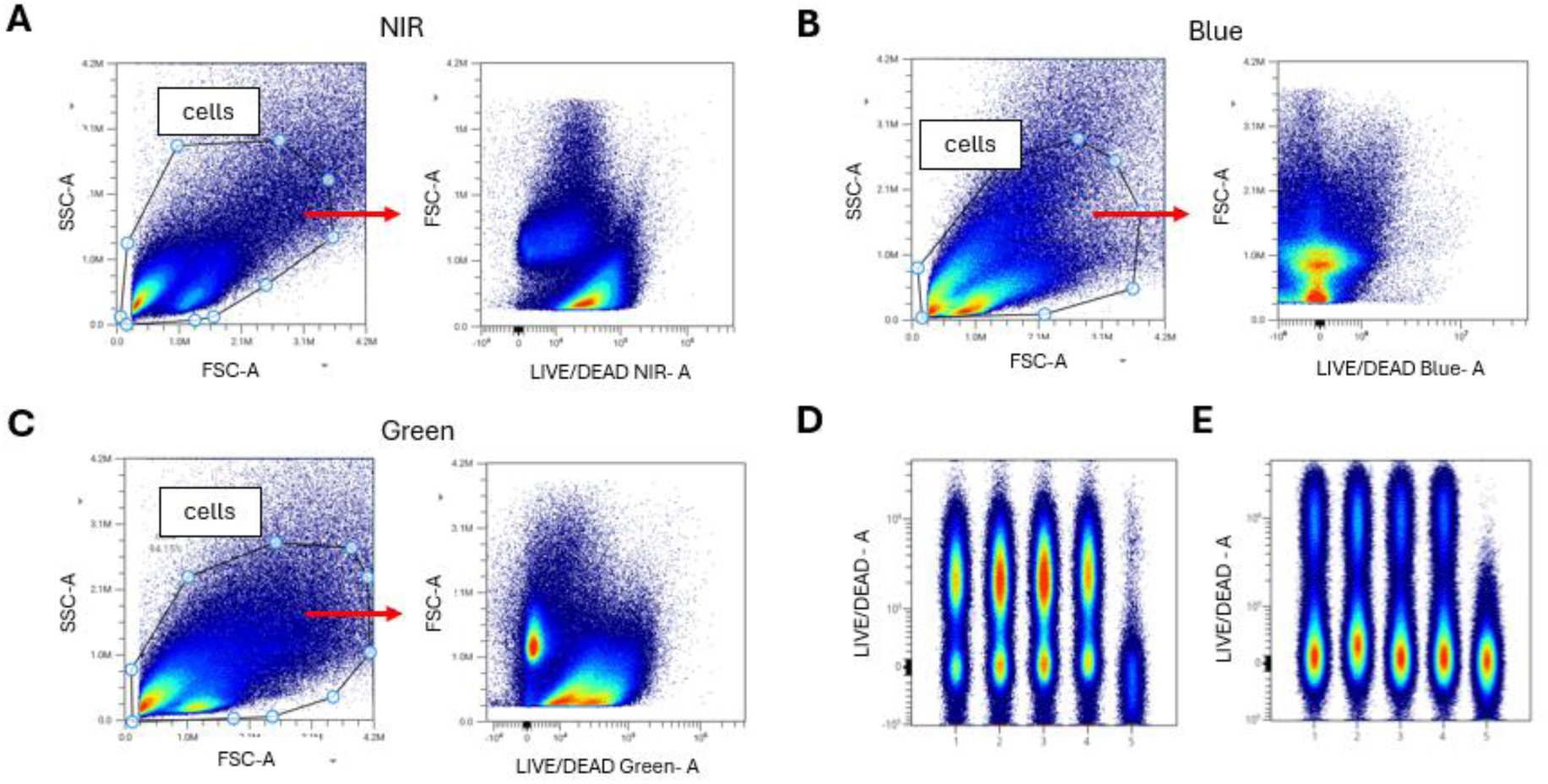
Optimisation of LIVE/DEAD Fixable dead cell staining. (A-C) Scatter plots of liver cell suspensions stained with (A) LIVE/DEAD NIR, (B) LIVE/DEAD Blue and (C) LIVE/DEAD Green Fixable dead cell dyes. (D, E) Titrations of the LIVE/DEAD Fixable dead cell dye on (D) heat killed and (E) non-heat killed cells from whole liver single cell suspensions. The dilutions used were (1) 1: 1,000, (2) 1: 2,000, (3) 1: 5,000, (4) 1: 10, 000 and (5) unstained.

For the viability dye titration experiments, heat-killed (90°C for 5 mins; Fig. 4D) and non-heat killed samples (Fig 4E) were stained separately. Before data acquisition, stained and unstained cells were mixed in the same tube, to obtain a clear positive and negative peak for each sample. The 1:5,000 dilution was the one with the highest Stain Index, providing the clearest resolution between live and dead cells with minimal signal distortion.

#### Antibody Selection

There are several factors that can influence the selection of conjugated antibodies for FSFC panels. It is essential to understand the properties of the target cells, particularly the expression levels of the chosen antigens and whether they are co-expressed on the same cell populations. Whenever possible, avoid combining antibodies that bind to epitopes located in close physical proximity, as this can impair antibody affinity through steric hindrance or stoichiometric inhibition.

When selecting fluorochromes, several considerations are important. Fluorochromes with significant spectral overlap or those contributing to unmixing-dependant spreading due to spillover, instrument noise and statistical variation, should ideally be assigned to antigens that are not co-expressed on the same cell. In addition, the gating strategy and co-expression of markers of interest should be considered when selecting fluorochromes. This approach helps to minimize data spread and improve the accuracy of spectral unmixing. Moreover, bright fluorochromes should be matched with antigens with unknown distribution and low expression levels, while dimmer fluorochromes are better suited for highly abundant targets with well-defined expression patterns. Using dim fluorochromes for low-abundance proteins can results in signal loss due to high background and AF. Similarly, assigning bright fluorochromes to highly expressed proteins may lead to excessive signal spread, which can compromize resolution and cause inaccurate spectral unmixing.

Availability of antibody clones and fluorochrome conjugates is also a major consideration. For non-immune cell markers, commercial demand is lower, which often limits the range of available conjugates and can constrain panel design. To determine the suitability of selected clones and fluorochromes for FSFC applications, we utilized databases such as <u>CiteAb</u> and <u>Biocompare</u>.

Titration of the selected antibodies (Table 1) was performed by incubation of individual antibodies over a dilution range from 1:20 - 1:640 in buffer containing 1:5,000 dilution of the viability dye. The optimal dilution for each antibody was then determined by assessing the separation of the positive and negative cell populations, with minimal background signal (Stain Index). For this assessment, only the live cells were taken into consideration, as dead cells can non-specifically bind to antibodies leading to high background and false positive signal. Finally, all dilutions were validated as part of a complete panel. The plots for all antibody titrations of this EC panel are presented in Fig. 5A-M, with red boxes highlighting the selected dilutions for each antibody.

**Figure 5:**
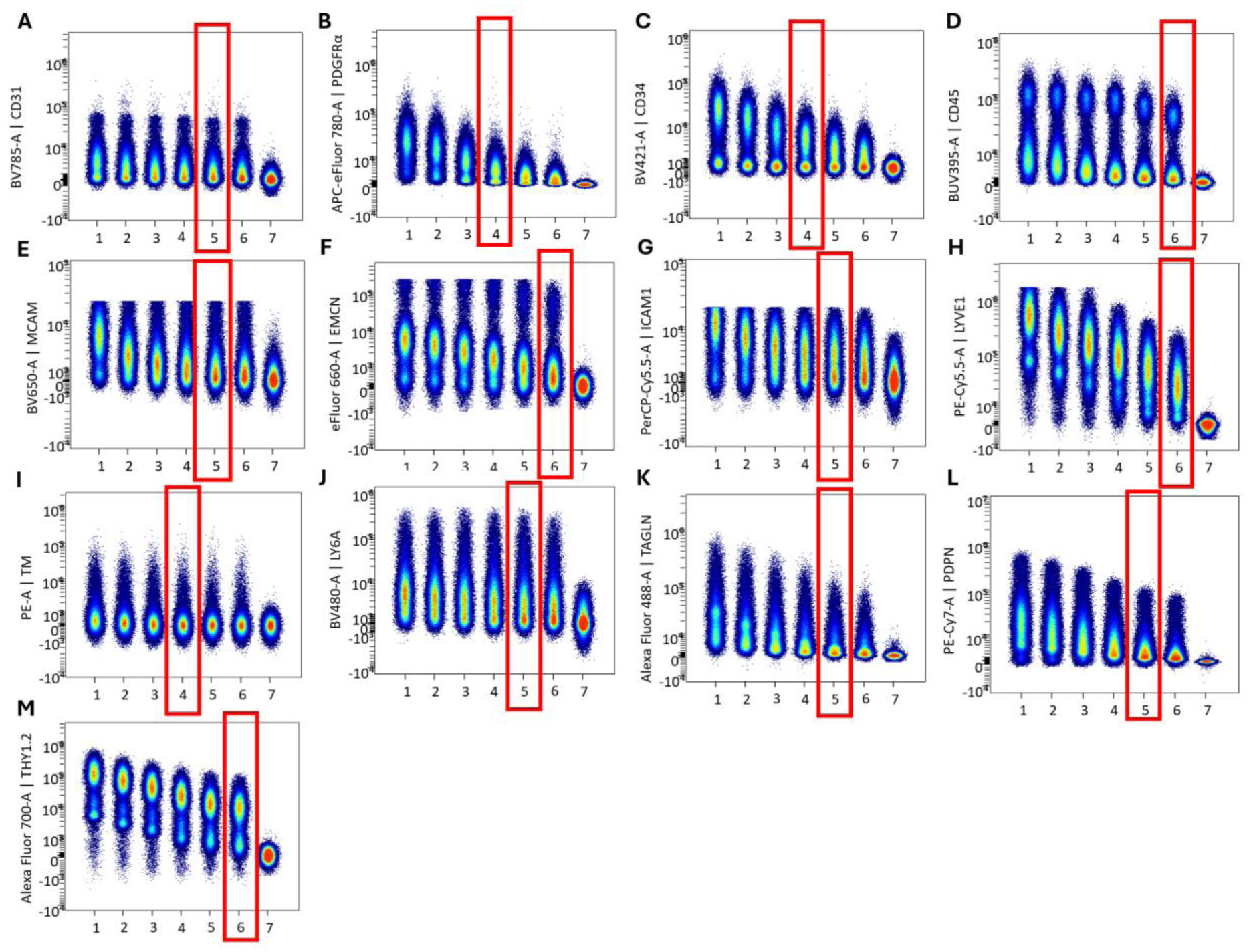
Titration of all endothelial cell panel antibodies. Each antibody was diluted at (1) 1:20, (2) 1:40, (3) 1:80, (4) 1:160, (5) 1:320, (6) 1:640, while an unstained sample (7) was also used for comparison. The antibodies tested included (A) CD31, (B) platelet-derived growth factor receptor α (PDGFRα), (C) CD34, (D) CD45, (E) melanoma cell adhesion molecule (MCAM), (F) endomucin (EMCN), (G) ICAM1, (H) LYVE1, (I) thrombomodulin (TM), (J) LY6A, (K) transgelin (TAGLN), (L) podoplanin (PDPN) and (M) THY1.2. Red boxes indicate the final dilution chosen for each marker.

**Table 1:** Key resources table.

| REAGENT or RESOURCE | SOURCE | IDENTIFIER |
| --- | --- | --- |
| **Antibodies** |  |  |
| CD31 (Clone 390) BV785 | Biologend | 102435 |
| CD45 (Clone 30-F11) BUV395 | BD Bioscience | 565613 |
| Thrombomodulin (Clone LS17-9) PE | Fisher Scientific | 12-1411-82 |
| EMCN (Clone V.7C7) eFluor660 | Fisher Scientific | 50-5851-82 |
| LYVE1 (Clone ALY7) PE-Cy5.5 | Novus | NBP1-43411 |
| PDPN (Clone 8.1.1) PE-Cy7 | Biologend | 127411 |
| MCAM (Clone ME-9F1) BV650 | Biologend | 740636 |
| Ly6A (Clone D7) BV480 | Biologend | 746663 |
| ICAM1 (Clone YN1/1.7.4) PerCP-Cy5.5 | Biologend | 116123 |
| TAGLN (Clone TAGLN/247) AF488 | Novus | NBP2-47688AF488 |
| Thy1.2 (Clone 30-H12) AF700 | Biologend | 105320 |
| PDGFRa (Clone APA5) APC-eFluor780 | Fisher Scientific | 47-1401-82 |
| CD34 (Clone MEC14.7) BV421 | Biologend | 119321 |
| **Chemicals, peptides, and recombinant proteins** |  |  |
| Phosphate Buffered Saline (PBS) | Fisher Scientific | BR0014G |
| Bovine Serum Albumin (BSA) | Fisher Scientific | BP9702-100 |
| EthyleneDiamineTetraacetic Acid (EDTA) | Sigma Aldrich | E7889-100ML |
| Collagenase | Gibco | 17018-029 |
| Dispase II | Roche | 04942078001 |
| DNase | Invitrogen | 18047-019 |
| PBS with CaCl<sub>2</sub> and MgCl<sub>2</sub> | Gibco | 14170-088 |
| 70µm cell sieve | Scientific Laboratory Supplies | 431751 |
| Fc blocking solution | Miltenyi Biotec | 130-092-575 |
| Live/Dead™ Green Fixable dye | Invitrogen | L23105 |
| Brilliant stain buffer | BD Biosciences | 554656 |
| Intracellular fixation and permeabilization buffer | Invitrogen | 00-5521-00 |
| Permeabilization wash buffer | Invitrogen | 00-8333-56 |
| Foetal Bovine Serum (FBS) | Gibco | A4766801 |
| UltraComp eBeads™ Compensation Beads | Fisher Scientific | 01-2222-41 |
| **Experimental models: Organisms/strains** |  |  |
| Carbon Tetrachloride CCl<sub>4</sub> | Sigma-Aldrich | 289116-100ML |
| Modified Clinton-Cybulsky cholesterol diet | IPS Product Supplies | 58R6 |
| Fructose | Sigma-Aldrich | F0127-1KG |
| Sucrose | Sigma-Aldrich | S9378-1KG |
| Peanut oil | Sigma | P2144-250ML |
| C57/BL6 Mice | Charles Rivers Labroatories |  |
| **Software and algorithms** |  |  |
| SpectroFlow® 3.3.0 software | Cytek® |  |
| Cytek® Cloud | Cytek® |  |
| OMIQ® | Dotmatics |  |
| Other |  |  |
| 4 laser Aurora (UV, V, B, R) spectral flow cytometer | Cytek® |  |

### Unmixing controls and extraction of complex autofluorescence signatures

The next step was to generate a library of single stain reference controls for all fluorochromes, as these represent an essential tool for correct data unmixing. Reference controls capture the unique spectral signature of each fluorochrome, enabling the software to correctly assign fluorescence signals to each fluorochrome across the spectrum. They should initially be tested on both beads and cells to determine the most accurate signal for each fluorochrome. For tandem dyes, the reference control should be prepared using the exact same antibody vial to minimize lot-to-lot variation. For non-tandem dyes, a different antibody may be used if necessary to achieve a strong positive signal and correct brightness. Examples of reference signatures for each fluorochrome can be found on the Cytek® cloud and in the Cytek® Aurora Fluorochrome Selection Guidelines. Additionally, the positive and negative signal for each reference control must originate from the same cell or bead population to ensure consistency. Beyond spectral signature accuracy, signal intensity is also critical: the reference control signal should be at least as bright as the fully stained sample; otherwise, unmixing errors will occur.

The unstained control is one of the most important and critical samples in spectral flow cytometry. It must consist of the exact same cell types as the fully stained samples and include sufficient cell numbers to represent all relevant subpopulations. This ensures accurate AF signatures extraction for each sample.

Solid tissue samples present additional complexity, as they contain a variety of different cell types, metabolites, extracellular matrix proteins and connective tissue. These factors contribute to unique AF signatures that vary with tissue composition and cellular complexity. Moreover, AF can be further influenced by chemicals, drugs, treatments, tissue injury, inflammation and fibrosis. Therefore, the most effective strategy for AF extraction in complex tissue and injury models is to capture all the different AF signatures to account for these variations. We used the ‘multi-AF extraction’ tool available through the SpectroFlo® software to identify and extract several AF signatures. It is crucial to include appropriate unstained control samples for all relevant variables, such as time-points, sex and injury models, to avoid potential unmixing errors caused by unique AF signals (Fig. 6 A-D).

**Figure 6:**
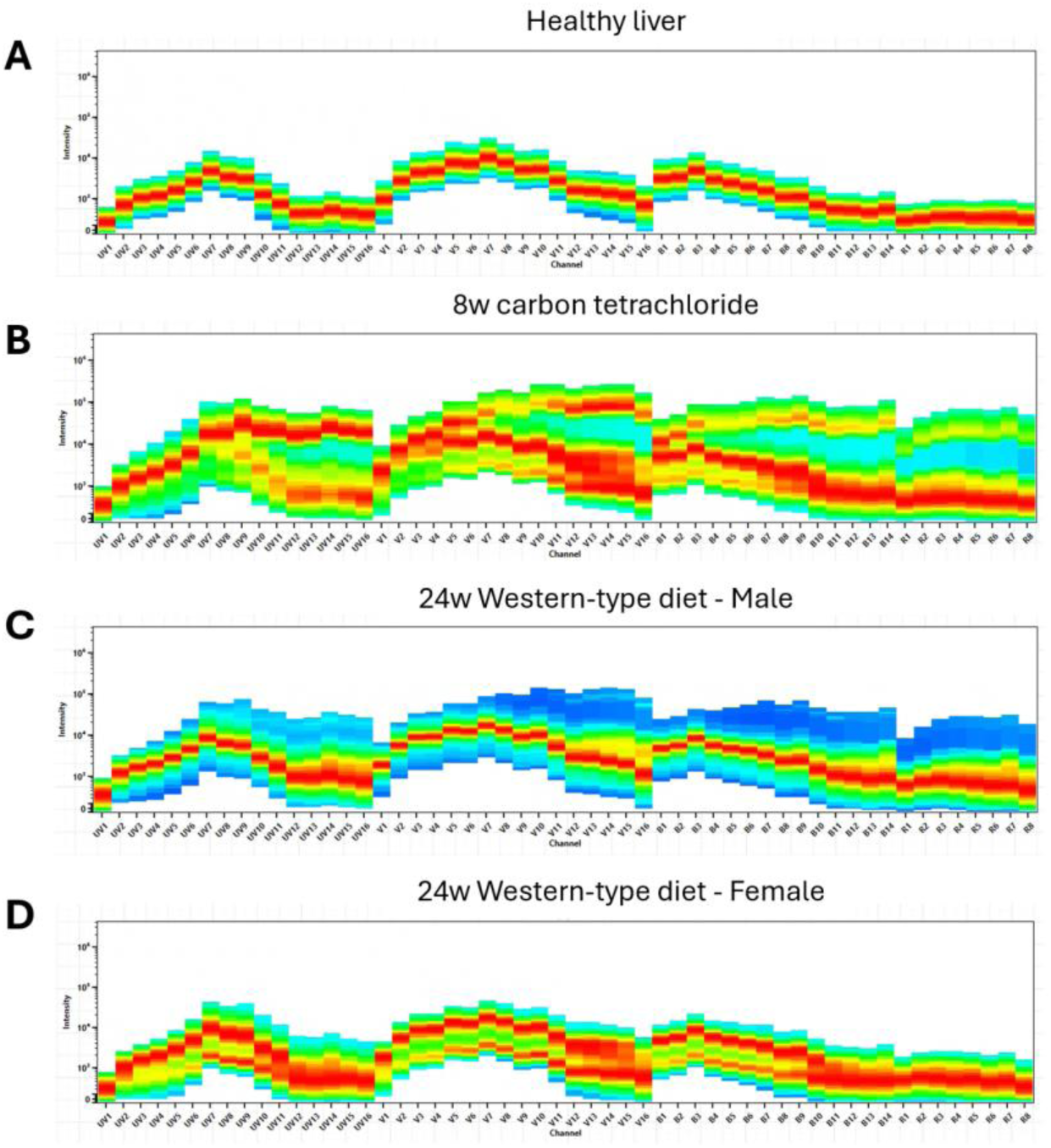
Examples of autofluorescence signatures in different murine liver samples. Spectral plots of unstained samples obtained from (A) healthy male liver tissue, (B) fibrotic liver tissue after 8 weeks of carbon tetrachloride injections in a male mouse, (C, D) steatotic liver after 24 weeks of a Western-type diet in (C) a male mouse and (D) a female mouse. The high diversity demonstrates the requirement for group-specific unmixing.

### Additional controls and data analysis

#### Fluorescence minus one (FMO) control samples

To establish a threshold for positive signal for each target, it is essential to include appropriate FMO controls in the experiment. As discussed previously, fluorochrome spread can increase background signal, making it difficult to distinguish true positive signal from noise. FMO controls address this by using all fluorochromes in the panel but one for each marker of interest. This approach provides a clear background threshold and defines the point beyond which a positive gate should be drawn for each fluorochrome (Fig. 7A-O).

**Figure 7:**
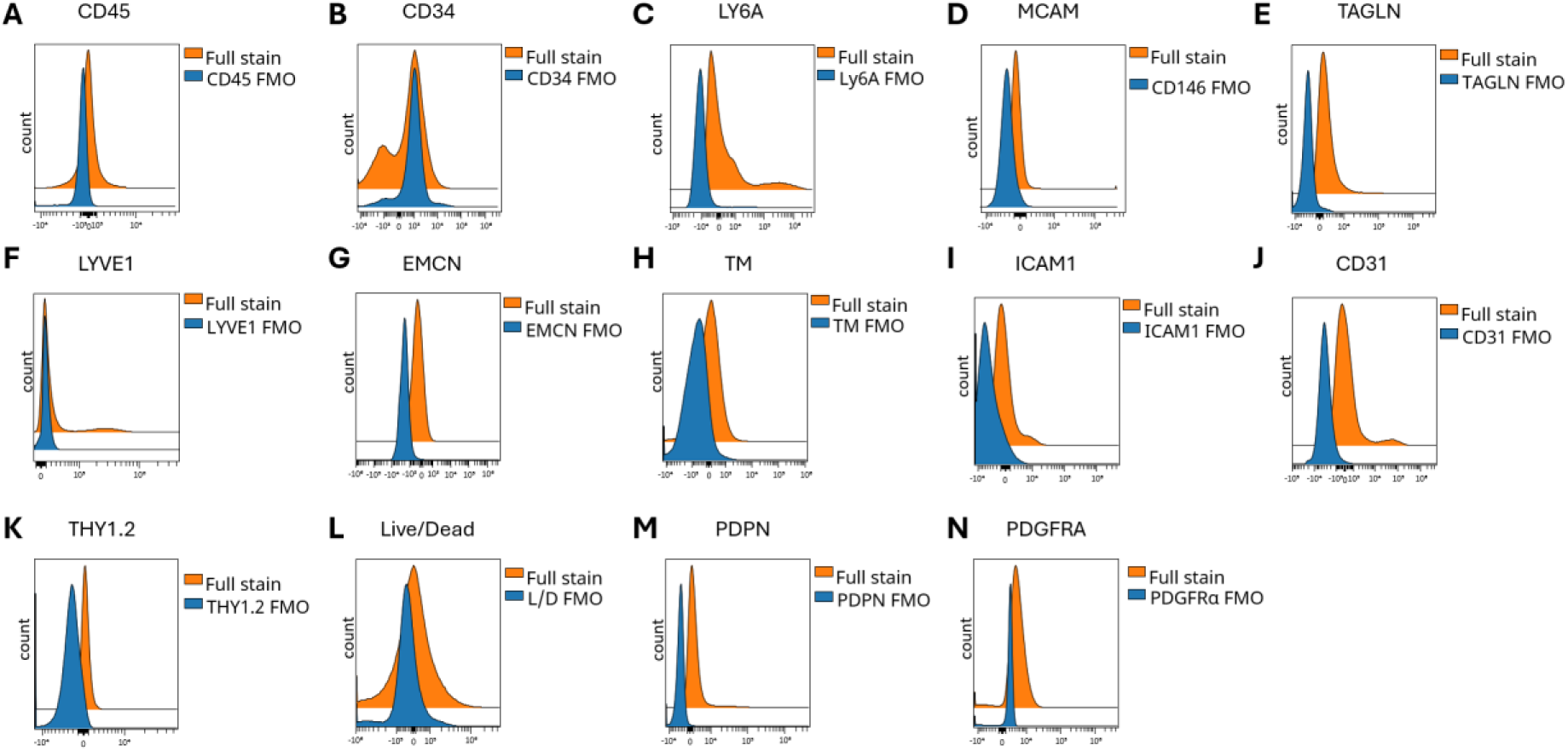
FMO controls for all markers of the EC panel. Histograms comparing Fluorescence Minus One (FMO; blue) samples to full stain samples (orange) in heathy liver cells for (A) CD45, (B) CD34, (C) LY6A, (D) melanoma cell adhesion molecule (MCAM), (E) transgelin (TAGLN), (F) LYVE1, (G) endomucin (EMCN), (H) thrombomodulin (TM), (I) ICAM1, (J) CD31, (K) THY1.2, (L) LIVE/DEAD Green Fixable dead cell dye, (M) podoplanin (PDPN), (N) platelet-derived growth factor receptor α (PDGFRα).

#### High dimensionality reduction

In addition to traditional gating analysis, FSFC allows for the use of dimensionality reduction analysis. Given the complexity of the panel and the diversity of cell phenotypes studied, clustering analysis can be used to identify cells sharing multiple properties, rather than relying on conventional manual NxN plots that analyse only two parameters at a time. Several software platforms support this type of analysis; however, for simplicity, we will describe our pipeline optimized within OMIQ® (Fig. 8).

**Figure 8:**
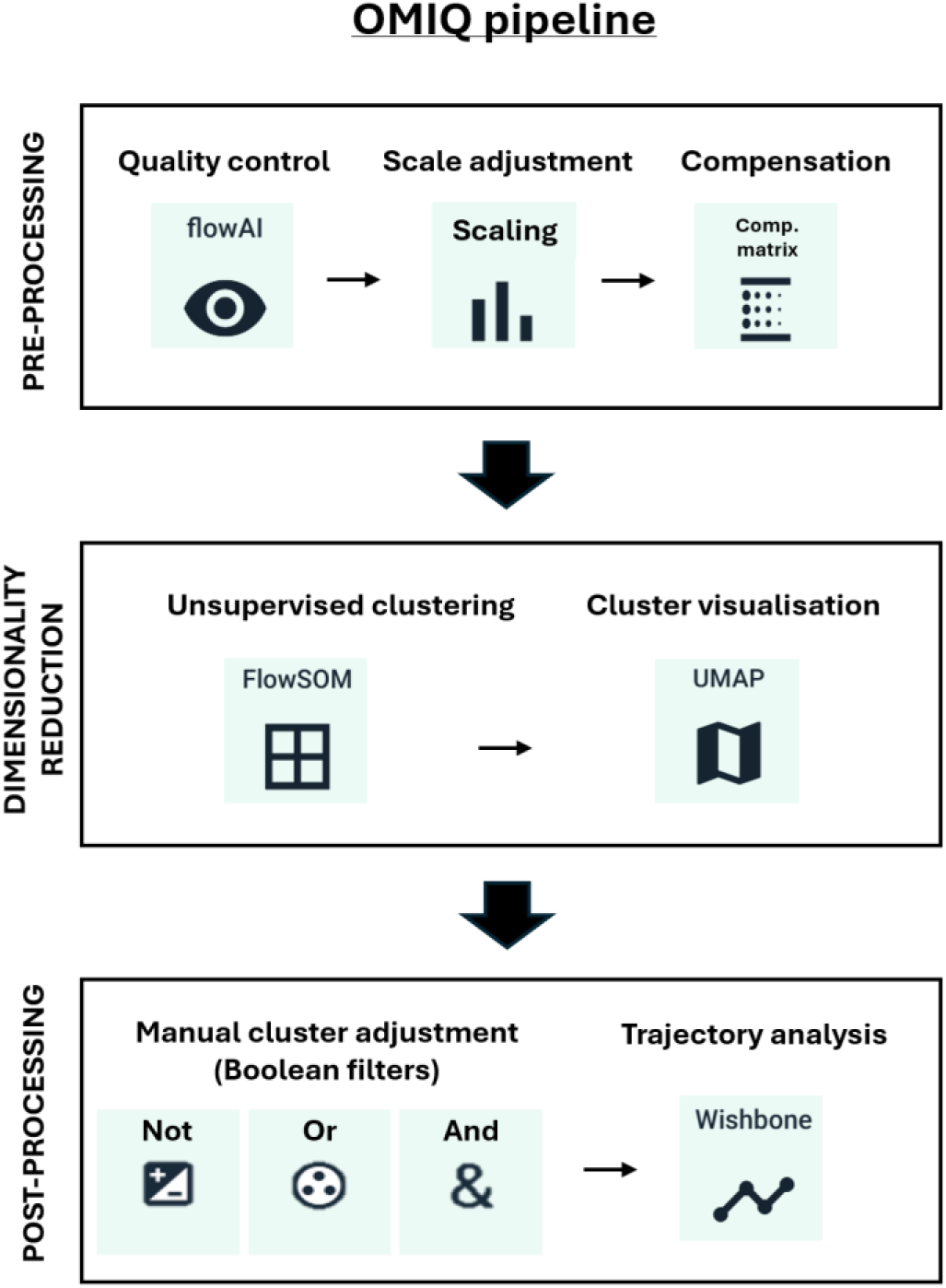
Outline of the OMIQ pipeline used for high dimensional analysis of full spectrum flow cytometry data. For data pre-processing, the FlowAI algorithm was used for quality control, followed by data scaling and manual compensation as required. Unsupervised clustering was then performed using the FlowSOM algorithm, while cluster visualization was achieved through UMAP plots. During post-processing, clusters were manually merged or split as deemed necessary, and trajectory analysis was conducted using the Wishbone algorithm.

Because signal acquisition can be affected by several factors such as the presence of air bubbles, laser and instrument fluctuation or changes in flow rate, a quality control (QC) step is essential to ensure data integrity. Various algorithms are available for QC; in our workflow we applied FlowAI ^17^ to identify and remove anomalous data acquired through variation in the flow rate and signal range over time. Importantly, uncompensated data should be used for this algorithm, allowing for unbiased QC, not affected by user-dependent manual compensation. FlowAI generates a new gate containing ‘clean’ data, which is then used for all downstream analyses.

After QC, data were properly scaled and compensated, before being manually cleaned to remove debris, doublets and dead cells (Fig. 3A-C). To identify unique EC subpopulations, we performed unsupervised clustering based on the 6 EC identity markers of the panel. Data from all experimental groups were concatenated prior to clustering, allowing subsequent analysis and comparison between samples groups. Clustering was undertaken using the FlowSOM ^18^ algorithm, which generates a self-organizing map that uses K nearest neighbours, where K is manually defined by the user. We selected K=20, anticipating more than 5 but fewer than 20 subpopulations (meta-clusters), based on previous scRNAseq studies ^13, 14^. Once clusters were generated, their properties were reviewed manually using the median fluorescence intensity (MFI) of their positive markers, and clusters were manually merged or split as needed using Boolean filters.

Dimensionality reduction plots can then be used to visualize the clusters generated. Depending on the data and biological questions, different types of plots can be used and the settings for map generation can be optimized to better present the data. Clustering data for hepatic and cardiac EC subpopulations (Fig. 9 A and B respectively) were visualized by Uniform Manifold Approximation and Projection (UMAP) plots ^19^. UMAP better preserves the global structure of the data allowing to clearly segregate each cluster and thereby visualize their expression profiles for each EC marker within the map (Fig. 8 C and D). Whilst the same EC panel can be applied across different tissues, clustering analysis must remain tissue-specific to accurately capture differences in marker expression profiles. For this reason, each UMAP visualisation was generated separately for each tissue type, and the UMAP plots cannot be directly overlaid. However, UMAP plots highlight the greater diversity of hepatic EC subpopulations compared to those in the heart, with 8 of the 12 hepatic clusters expressing more than 4 EC identity markers, compared to only 1 of 7 within the heart. Additionally, the composition of heart EC is dominated by a single, large subpopulation while hepatic sub-populations are more evenly distributed between the clusters (Fig. 9E).

**Figure 9:**
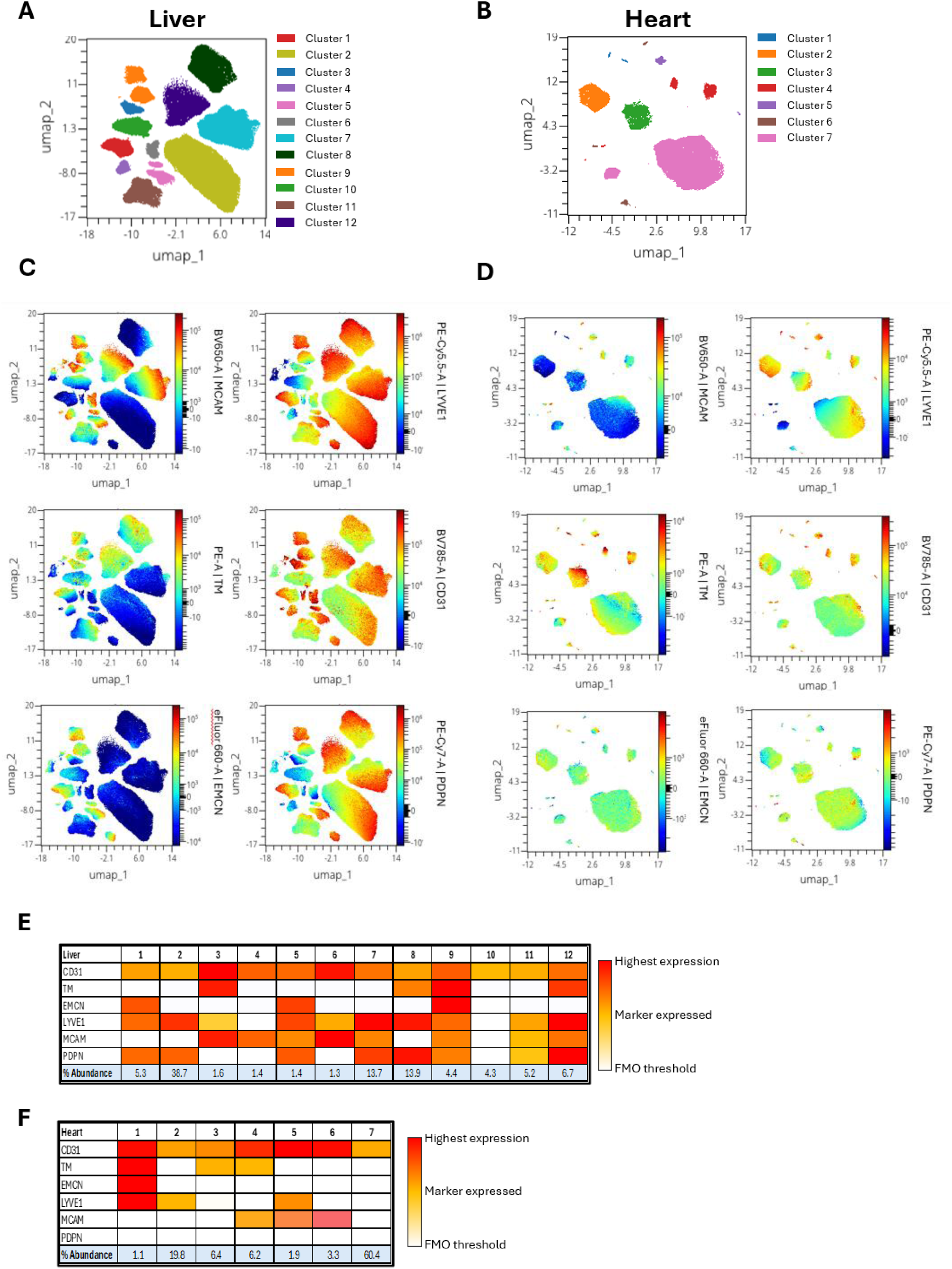
Dimensionality Reduction analysis of hepatic and cardiac endothelial cells. (A, B) UMAP plots generated using the endothelial markers CD31, thrombomodulin (TM), endomucin (EMCN), LYVE1, melanoma cell adhesion molecule (MCAM), and podoplanin (PDPN) for CD31^+^CD45^-^ endothelial cells from healthy murine (A) liver and (B) heart tissue. (C, D) Expression of each marker was overlaid onto the UMAP to distinguish the characteristics of each cluster for (C) liver and (D) heart samples. (E, F) Heatmaps of the mean fluorescence intensity of each EC identity marker used to define the 12 hepatic EC clusters (E) and the 7 cardiac EC clusters (F). The colour denotes the intensity of expression of a marker; below the FMO threshold (white), expression above the FMO threshold (yellow) and highest expression (red). The abundance of each cluster expressed as a percentage of the total EC is also presented.

#### Application of secondary analytical tools to EC sub-populations

As our experimental approach included multiple timepoints and measured markers of activation status, we had the opportunity to assess cellular dynamics by modelling their trajectory. This was achieved by applying Wishbone trajectory analysis which has previously been published to infer T-cell differentiation ^20^. Wishbone enables an unbiased assessment of cell plasticity from a manually identified starting cell population. In this example we began with EC that had low activation status defined as ICAM1^-^ TAGLN^-^ (Fig. 10A). To determine the trajectory of activation of these cells we used Wishbone analysis which identified a trunk and two branches (Fig. 10B) based on the changes in expression of ICAM1, LY6A, CD34, TAGLN, THY1.2 and PDGFRα. The Wishbone algorithm allocates ‘milestones’ for each cell analysed, building a trajectory that can be linear, or include a bifurcation point^21^, providing insight into lineage relationships^10^. These criteria enabled us to visualize the phenotypic changes that occur time in response to Western diet (Fig. 10C).

**Figure 10:**
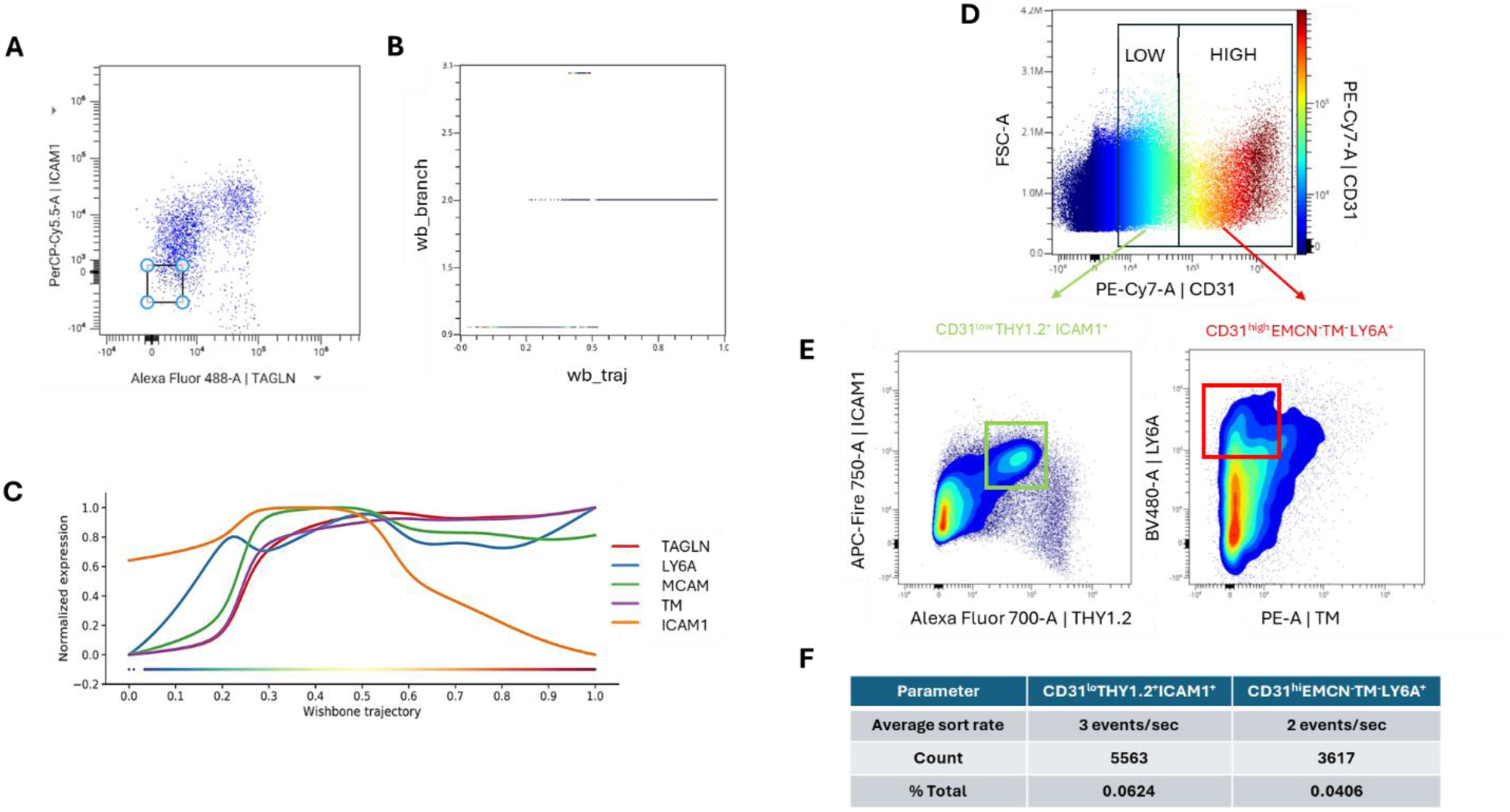
Secondary analysis of EC subpopulations by Wishbone trajectory and full spectrum cell sorting. (A-C) Example workflow for generating a Wishbone trajectory plot. (A) Dot plot of TAGLN vs ICAM1 to identify the starting gate for Wishbone analysis, defined as TAGLN^-^ ICAM1^-^ cells. (B) Branch and trajectory plot which allocates milestones for each cell analysed. (C) Wishbone trajectory plot for TAGLN, LY6A, MCAM, TM and ICAM1 highlighting the transition of EC in response to 12w Western diet. (D) Dot plot of CD31 expression in whole liver cell suspension from mice treated with carbon tetrachloride for 4 weeks. (E) Full spectrum cell sorting of CD31^low^ THY1.2^+^ ICAM1^+^ (left; green gate) and CD31^high^ TM^-^LY6A^+^ (right; red gate). (F) Table of sorting efficiency parameters for each population.

Using our initial EC panel, we demonstrated changes in inflammatory status and induction of Endothelial-to-Mesenchymal Transition (EndMT) associated with murine models of progressive liver fibrosis^10^. This comprehensive phenotyping approach allowed the isolation of rare and complex EC subpopulations through full spectrum cell sorting. We identified a pro-fibrotic EC phenotype, expressing the EndMT marker THY1.2, which was preferentially expressed on CD31^low^ ECs. In contrast, a LY6A^+^, pro-inflammatory EC phenotype that was preferentially acquired by CD31^high^ ECs (Fig. 10D and E) after 4 weeks of carbon tetrachloride (CCl_4_)-induced liver fibrosis. These findings informed the development of a targeted cell-sorting panel which was transferable to the Cytek® Aurora CS cell sorter system (Table 2) with the help of Cytek®. While antibody clones remained unchanged, the panel was streamlined to reduce complexity and improve separation, thereby enhancing sort efficiency and resolution of these rare EC subpopulations (Fig. 10F).

**Table 2:** Antibody panel for full spectrum cell sorting

| REAGENT or RESOURCE | SOURCE | IDENTIFIER |
| --- | --- | --- |
| Antibodies |  |  |
| CD31 (Clone 390) PE-Cy7 | Biolegend | 102417 |
| CD45 (Clone 30-F11) BUV395 | BD Bioscience | 565613 |
| Thrombomodulin (Clone LS17-9) PE | Fisher Scientific | 12-1411-82 |
| EMCN (Clone V.7C7) eFluor660 | Fisher Scientific | 50-5851-82 |
| LYVE1 (Clone ALY7) PE-Cy5.5 | Novus | NBP1-43411 |
| Ly6A (Clone D7) BV480 | Biolegend | 746663 |
| ICAM1 (Clone YN1/1.7.4) PerCP-Cy5.5 | Biolegend | 116123 |
| Thy1.2 (Clone 30-H12) AF700 | Biolegend | 105320 |
| CD34 (Clone MEC14.7) BV421 | Biolegend | 119321 |

## Discussion

Flow cytometry techniques have provided remarkable insight into the identity and phenotype of circulating cell types, such as leukocytes ^22^, platelets ^23^ and cancer cells ^24^, and have become a gold standard technique in immuno-oncology ^2^. The iterative process of acquiring and identifying new cellular phenotypes has driven the emergence of new technologies, such as FSFC, capable of resolving more than 50 fluorochromes ^25^. However, applying these techniques to solid tissue-derived cells has been limited by several challenges, including tissue dissociation, complex tissue- and cell-specific AF signatures and availability of appropriate antibody reagents. To address this knowledge and application gap, we established a 14-colour EC panel designed to identify specific EC subpopulations and assess phenotypic changes induced in response to tissue fibrosis. Although ECs are highly diverse, as shown in numerous scRNAseq studies ^4, 5, 6, 26^, proteomic phenotyping remains comparatively underexplored. We aimed to integrate our knowledge of vascular heterogeneity in health and disease to identify key markers of EC identity, inflammation and fibrosis using commercially available antibodies to develop a framework for an EC-specific FSFC panel.

ECs represent a particularly challenging population for flow cytometry due to their high and complex AF signatures ^7^. We addressed these hurdles by incorporating a multitude of controls to identify confounding factors that influence spectral unmixing. Notably, healthy livers displayed between 3 and 4 AF signatures, whereas mice treated for 8 weeks with CCl_4_ had up to 7 distinct AF signatures, significantly increasing the complexity score of the panel. Initially, model- and time-point-specific AF signatures required laborious manual identification and removal. However, the release of the Cytek® Multi-AF extraction tool in 2024 has dramatically simplified multi-AF identification and improved its removal within an unmixing workflow.

Our core EC identity markers displayed remarkable diversity across the EC sub-populations, enabling the resolution of 12 EC subpopulations in the liver and 7 within the heart. These are diverse markers of EC identity and are associated with a variety of different functions^27^. While direct comparison between liver and heart EC populations is not feasible, we observed clear tissue-specific demographics. For example, LYVE1 and PDPN expression was far more prominent in the liver compared to the heart, reflecting the specialized nature of hepatic ECs, which share traits with lymphatic vessels ^28^. Furthermore, by incorporating activation markers, we could distinguish phenotypic changes occurring in the liver in response to a model of liver fibrosis associated with inflammatory status and fibrosis-induced EndMT (these findings are described in detail in Gkantsinikoudi *et al.* 2026^10^). We also show that Wishbone analysis provides novel insights into EC plasticity. Finally, we demonstrate that these unique characteristics can be leveraged for full spectrum cell sorting, enabling isolation of rare EC subpopulations for further downstream analysis.

In summary, we demonstrate the successful application of an EC-specific FSFC panel to liver and heart tissues and its utility in models of liver injury. This work establishes a robust methodological foundation that will accelerate high-dimensional phenotyping of complex cell populations and expand the capabilities of FSFC in solid tissue research.

## Acknowledgement

Dr. Stephan Kaizik and Dr. Adam Davison at Cytek Biosciences, Cambridgeshire, CB6 2HF, United Kingdom for technical support and use of the Aurora CS Sorter and the Flow Cytometry Core Facility, Babraham Institute, Cambridge, CB22 3AT, United Kingdom. This research was supported by Dr Paul Imbert, manager of the CMR Advanced Bioimaging Facility (CMR-ABF) and the Barts Cancer Institute (BCI) / William Harvey Research Institute (WHRI) Flow Cytometry Facility which are located in the John Vane Science Centre, Charterhouse Square, Queen Mary University of London, EC1M 6BQ, United Kingdom.

## Funding

This project was funded by Medical Research Council, UK (MR/Y013751/1; NPD), British Heart Foundation Doctoral Training Programme, UK (FS/4yPhD/F/21/34161; CG) and Bart’s Charity London, UK (MGU0514; NPD).

## Motivation

Single-cell RNA sequencing has transformed our understanding of cellular heterogeneity, but transcriptomic data alone cannot capture information regarding protein expression, post-transcriptional modifications or cell-surface interactions. Full spectrum flow cytometry (FSFC) offers high-dimensional proteomic analysis at single-cell resolution, yet its application to non-immune cells, particularly endothelial cells (EC) from solid tissues, is limited due to their complex autofluorescence signatures. We developed and optimized a FSFC protocol to overcome this challenge, enabling robust phenotyping of endothelial subpopulations, downstream high dimensional analysis and isolation of rare EC subpopulations from murine liver and heart tissues.

## Limitations of the study

The FSFC protocol described has several inherent limitations. As experiment-specific AF controls are required for accurate unmixing, variability between datasets can be introduced, necessitating larger biological sample numbers to achieve robust and reproducible results. Moreover, unlike scRNAseq, which enables unbiased discovery of transcriptional states, FSFC-based cluster identification is constrained by the markers included in the antibody panel. The limited choice of fluorochrome conjugates of non-immune cell targets is an important challenge that if addressed, could unlock the discovery of rare or unexpected cell phenotypes.

## EXPERIMENTAL MODELS

### Animals

All procedures used in this study were approved by the Animal Use and Care Committee of Queen Mary University of London (QMUL) and were in accordance with national and international regulations. C57BL/6 mice at 8-10 weeks old were used for all experiments. Food and water were provided ad libitum. Individually ventilated cages were used in a temperature-controlled facility. A 12-hour light: 12-hour dark cycle was maintained.

### Induction of liver injury using CCl_4_ injections

CCl_4_ was diluted 1 in 5 in peanut oil and was administered with an intraperitoneal injection at a dose of 0.8mg/Kg in 100μl. Injections were administered twice a week 4 or 8 weeks. No treatment or sham injections of 100μl peanut oil were used as a control.

### Induction of metabolic dysfunction-associated steatohepatitis using a Western diet

To model a Western diet and induce hepatic steatosis and fibrosis, mice were fed the modified Clinton-Cybulsky cholesterol diet up to 24 weeks. The water was also supplemented with carbohydrates (42g/L) consisting of 55% fructose and 45% sucrose.

## METHOD DETAILS

### Isolation of mouse liver and heart EC

1. Euthanize mice by cervical dislocation and exsanguinate by severing the carotid artery.

a. Perfuse with sterile deionized phosphate buffered saline (PBS) with 0.5% bovine serum albumin (BSA) and 2mM EDTA through the left ventricle.
b. Collect the heart and liver into ice cold PBS.
2. Dissociate tissues in the lab.

a. Weigh tissues before dissociation.
b. Mechanically disrupt tissues until homogenized using scissors.
c. Perform enzymatic digestion using 3ml pre-warmed (37°C) digestion buffer containing 2mg/ml collagenase, 8U/ml dispase II and 50U/ml DNase dissolved in PBS with CaCl_2_ and MgCl_2_.
d. Incubate at 37°C for 30 minutes on a shaker at approximately 180 rotations per minute (rpm). Note: Tissue weights can be used to calculate cells acquired per gram allowing normalization of cell number between samples. CRITICAL: The presence of Ca^2+^ and Mg^2+^ ions in digestion buffer are essential for activation of enzymes while DNase reduces cell clumping and therefore reduces the presence of cell doublets in FSFC.
3. Remove undissociated and connective tissue to obtain single cell suspension.

a. Filter homogenates through 70μm cell sieves.
b. Use a 2.5ml syringe plunger to gently dissociate any remaining tissue through the sieve.
c. Wash the sample tube with 1ml Wash Buffer (PBS with 0.5% BSA and 2mM EDTA) and pass through the sieve.
d. Wash the sieve with a further 1ml Wash Buffer.
e. Centrifuge cell suspensions for 5 minutes at 400g at 4°C and remove supernatant before resuspending in 1ml Wash Buffer.
f. Decant cell suspension through a fresh 70μm cell sieve and wash through with a further 1ml Wash Buffer.
g. Centrifuge cell suspensions for 5 minutes at 400g at 4°C and remove supernatant before resuspending in appropriate volume of Wash buffer for antibody staining. Note: We suggest resuspending the cell pellets in 200μl for profiling EC of the heart and 650μl for liver EC profiling. CRITICAL: Choose a resuspension volume that allows for additional controls to be run including unstained and Fluorescence Minus One (FMO) controls.

## Surface staining

1. Transfer 50μl of cell suspension to each well of a V-bottomed 96-well plate.

a. Wash with 200μl PBS by centrifuging the samples at 400g for 5 minutes.
b. Resuspend cell pellets in 50μl Fc blocking solution (diluted 1:500 in PBS) containing the LIVE/DEAD Fixable dead cell dye.
c. Incubate plate on ice for 30 min in the dark.
d. Wash with 200μl PBS by centrifuging the samples at 400g for 5 minutes at 4°C.
2. Prepare antibody staining panels ensuring you have the following groups:

a. Complete panel containing optimized dilutions of each individual antibody diluted in the Brilliant Stain Buffer.
b. Fluorescence FMO control samples from which each individual antibody is excluded in turn.
c. Unstained control samples. Example: A 5-stain panel containing antibodies A, B, C, D and E would require 7 wells for each sample assessed:

1: A, B, C,D, E (Complete panel)
2: B, C, D, E (FMO for A)
3: A, C, D, E (FMO for B)
4: A, B, D, E (FMO for C)
5: A, B, C, E (FMO for D)
6: A, B, C, D (FMO for E)
7: Antibody diluent only (Unstained control) Note: Staining performed in 5ml FACS tubes can allow for higher volume for washes steps; for staining performed in plates, leave an empty well around each sample to avoid cross contamination. Additional washes may be required for complex tissue digests.
3. Antibody panel staining.

a. Resuspend cells in 50μl of each antibody panel to individual wells.
b. Incubate plate on ice for 30 min in the dark.
c. Wash 2x with 200μl PBS by centrifuging the samples at 400g for 5 minutes at 4°C.
4. (Optional) Intracellular Staining. (Required for TAGLN staining in our panel).

a. Resuspend cells in 50μl the intracellular fixation and permeabilisation buffer.
b. Incubate plate at room temperature for 20 min in the dark.
c. Wash with 200μl permeabilization wash buffer by centrifuging the samples at 400g for 5 minutes.
d. Add 50μl of each antibody dilution to individual wells (diluted in permeabilisation wash buffer).
e. Incubate plate at room temperature for 30 min in the dark.
f. Wash x2 with 200μl permeabilisation wash buffer by centrifuging the samples at 400g for 5 minutes.
g. Resuspend the samples in 300μl MACS buffer (PBS with 2% FBS, 2mM EDTA).
h. Transfer samples to FACS tubes.

## Protocol for single stain reference controls

The staining protocol described above can be used to create single cell reference controls using cells. For staining of compensation beads to be used as single cell reference controls details are below:

a. Vortex beads.
b. Add 20μl beads in each well along with 80μl PBS (beads quantity and volume can be adjusted).
c. Add 1μl antibody per well (antibody amount based on titration).
d. Incubate plate at 4°C for 30 min in the dark.
e. Wash with 200μl MACS buffer by centrifuging the samples at 400g for 5 minutes.
f. Remove the supernatant.
g. Resuspend beads in 200μl MACS buffer.
h. Transfer samples to FACS tubes.

CRITICAL: When setting up these controls, it is important to initially stain both cells and beads to determine which provides the most accurate signal profile for each fluorochrome. Beads often offer a clear, fluorochrome-specific signature that does not rely on cellular expression, which is particularly useful when antibodies target low-abundance markers or rare populations. However, in some cases, the spectral profile of a fluorochrome bound to beads does not fully match its profile when bound to cells, potentially leading to spectral unmixing errors. Therefore, testing both options is essential to ensure optimal unmixing accuracy.

## Notes

### Competing Interest Statement

The authors have declared no competing interest.

### Summary of Updates

Addition of in a new reference throughout the text that is important as linked submission

